# A shift in developmental allometry underlies the transition to a multi-ovulate strategy from a single-ovulate ancestral state in *Phlox* (Polemoniaceae)

**DOI:** 10.64898/2026.01.06.697797

**Authors:** Bridget Bickner, Elena M Kramer

**Affiliations:** Department of Organismic and Evolutionary Biology, Harvard University; Arnold Arboretum of Harvard University; Department of Botany and Plant Sciences, University of California, Riverside

**Keywords:** *Phlox*, ovule primordium, seed number, allometry, reproductive fitness

## Abstract

The number of seeds produced per flower is influenced by many pre- and post-pollination factors and varies enormously across taxa, from one to thousands. This diversity has obvious implications for fitness and often reflects a tradeoff between investment in allocation per seed vs. the number of seeds produced. However, we know remarkably little about the developmental basis for variation in seed number, which, at its origin, is fundamentally variation in ovule number. In this study, we have used between and within species variation in the genus *Phlox* to explore the developmental basis of a shift between an ancestral state of producing only one ovule per locule and a derived state of producing two or more ovules per locule. Our results demonstrate that a change in the allometric relationships between ovary size, ovule size and ovule number has repeatedly evolved in this genus. This is the first study to demonstrate a relationship between ovule primordia size and ovary size and the first to find that the relative size of ovule primordia correlates with ovule number variation.

## INTRODUCTION

Seed size and number are fundamental measures of maternal reproductive fitness in plants. While total seed number depends on factors such as flower number and pollen deposition (reviewed in Primack 1987), ovule number per flower provides the starting point for expected seed yield per fruit. A tradeoff between seed size and number has been widely reported across all taxonomic scales: within species, between closely-related lineages, and across major plant lineages (Greene and Johnson 1994; Turnbull et al. 1999; Jakobsson and Eriksson 2000; Greenway and Harder 2007; Bawa et al. 2019). Previous work has highlighted that quantitative variation in ovule number can be produced by the same genotypes across different environments (plasticity) or by a variety of genetic mechanisms that usually have highly pleiotropic effects on ovary morphology (Brothers et al. 2016; Yuan and Kessler 2019; Cucinotta et al. 2020; Nepal et al. 2023). However, we know surprisingly little about the developmental basis of variation in the number of ovules per ovary (hereafter termed “ovule packaging strategy”, consistent with (Burd 1995).

Ovule development inside the ovary begins with the formation of primordia on the surface of meristematic tissue known as the placenta, and the number of primordia that form depends on multiple factors such as the size of the ovule primordia, the spacing of the primordia, and the available surface area of placental tissue (Cucinotta et al. 2020; Kawamoto et al. 2020; Shivaprakash and Bawa 2022). Although alteration of the spacing between primordia and the surface area of the placenta have both been demonstrated to produce ovule number variation (reviewed Cucinotta et al. 2020), ovule primordium size has received surprisingly little attention. As with many morphological traits, changes in the size of one structure often have pleiotropic effects on the size or number of related structures. For example, in many multi-ovulate species, an increase in ovary size also increases placenta surface area, resulting in an increase in the number of ovule primordia (Cucinotta et al. 2020; Qadir et al. 2022). However, maintaining allometric relationships can constrain the trait combinations upon which selection may act (Wessinger and Hileman 2016; Kalisz and Kramer). Much of our current understanding of variation in ovule number comes from lineages with a fixed multi-ovulate packaging strategy and a stable allometric relationship, particularly *Arabidopsis thaliana* or *Brassica napus*, in which ovule number changes quantitatively with ovary size and without major shifts in trait correlations (Yuan and Kessler 2019; Jiang et al. 2020; Qadir et al. 2022).

To investigate how major evolutionary transitions in ovule packaging strategy occur and what their resulting effect is on ovary and ovule (and fruit and seed) morphology, we leveraged the exceptional natural variation found in the genus *Phlox*. Most species of *Phlox* produce three seeds per fruit (one seed per locule in a three chambered-ovary); however, six species scattered across the genus (and representing each of the five major clades) appear to have independently evolved a multi-ovulate phenotype with at least two ovules per locule (Fig. 1; Gray 1870; Wherry 1955). Although a fully resolved phylogeny for *Phlox* is not available, existing phylogenetic evidence suggests that most, if not all, of the six species represent independent origins of the multi-ovulate strategy from a uni-ovulate ancestor (Ferguson and Jansen 2002; Landis et al. 2018; Garner et al. 2024). This transition from single-ovulate to multi-ovulate allows at least twice as many seeds to be produced per flower. Five species (*P. nivalis, P. buckleyi, P. longifolia, P. nana* and *P. sibirica* - the last of these not discussed in the present study) produce one, two, three, or rarely four ovules per locule (Gray 1870; Wherry 1930; Wherry 1955; B. Bickner, pers. obvs.), while one species, *P. roemeriana*, produces four to eleven ovules per locule (Ott et al. 1998; Lendvai and Levin 2003). Thus, the evolution of the multi-ovulate strategy represents both a shift in ovule packaging but also a transition from a fixed number of ovules per locule to a plastic number of ovules. In this study, we aim to (1) describe the morphology of early ovule development across *Phlox* ovule packaging morphs, (2) test the hypothesis that a tradeoff between seed size and number results in smaller seeds in multi-ovulate species, and (3) characterize changes in the allometric relationship between ovary/fruit and ovule/seed size among ovule packaging morphs and across developmental time. Collectively, our results demonstrate that a distinct change in the allometric relationships between ovary size, ovule size and ovule number has repeatedly evolved in this genus, highlighting the importance of carefully considering development in any study of seed number.

**Fig. 1.**
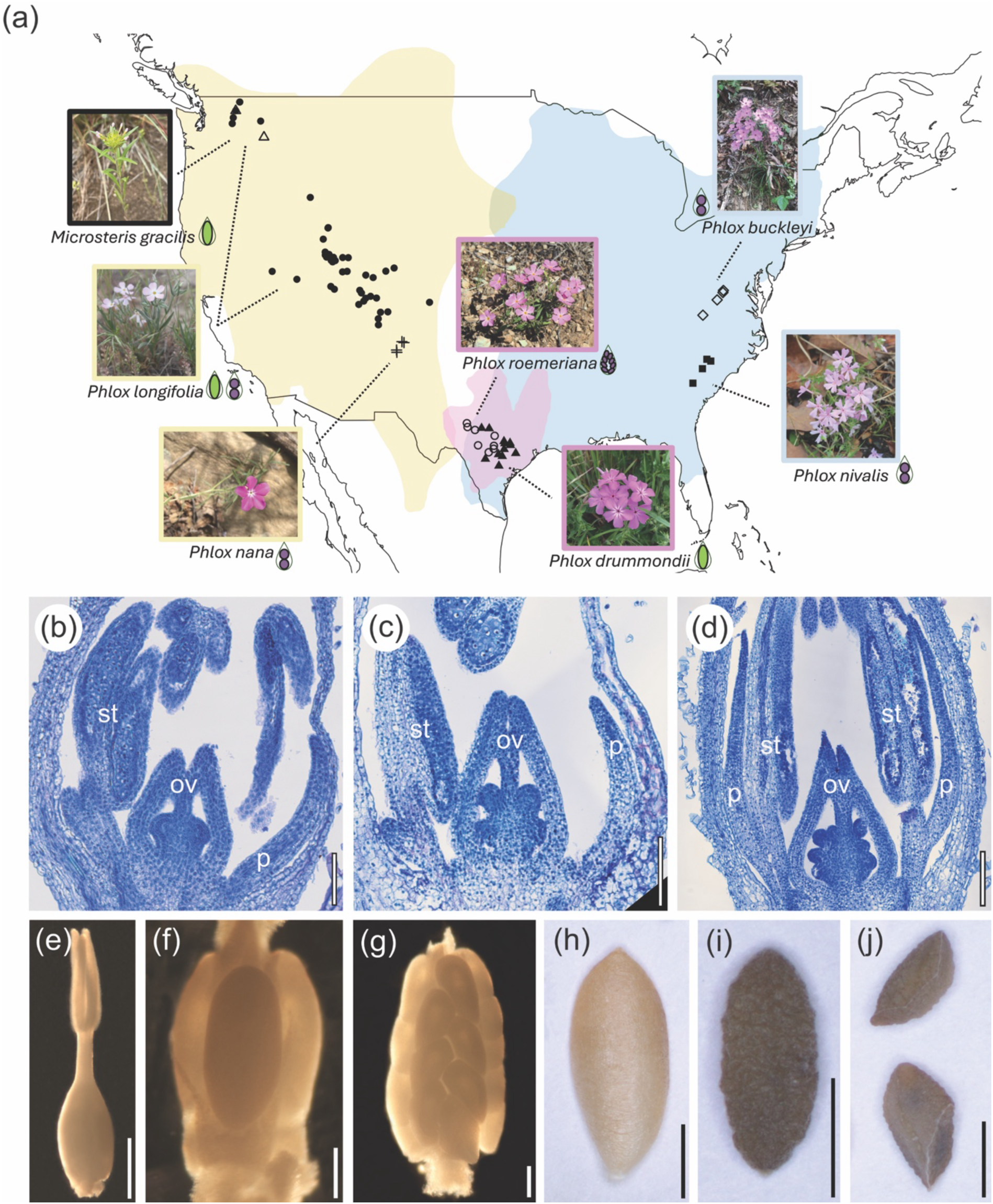
Geographic distribution of study taxa with exemplar dissected and histological sample materials. (a) Geographic distribution of the seven study taxa with the locations of sampled populations: the single-ovulate *Microsteris gracilis* (empty triangles) and *Phlox drummondii* (filled triangles) and the multi-ovulate *P. longifolia* (filled circles), *P. nana* (crosses), *P. nivalis* (filled squares), *P. buckleyi* (empty diamonds) and *P. roemeriana* (empty circles). Note that the overlap of two populations of *M. gracilis* and *P. longifolia* creates the appearance of a filled triangle in the northwest corner of the map. Each species is also designated by a schematic indicating the green single-ovulate and purple multi-ovulate conditions. Distributions are based on Wherry (1955) and Hopkins et al. (2014). The colored frame around each photograph indicates the phylogenetic classification of the species: black, outgroup representative; beige, western phlox; pink, Texas annual phlox; and blue, eastern standing phlox (Wherry 1955; Landis et. al. 2018; Ferguson et. al. 2002; Garner et al., 2024). (b-d) Histological sections of *P. drummondii* (b), *P. buckleyi* (c), and *P. roemeriana* (d). Organs are labeled as p=petal, st=stamen, ov=ovary. (e-g) Light micrographs of fixed material: pre-pollination pistil (e) and ovule (f) of *P. drummondii*, and ovules of *P. roemeriana* (g). (h-j) Light micrographs of mature fruit valve (h), fixed single-ovulate seed (i) and multi-ovulate seeds (j) of *P. longifolia*. Size bars in (b-d,f-g) = 200 µm; (e) = 500 µm; (h-j) = 2000 µm.

## METHODS

### Study system

*Phlox* (family Polemoniaceae) is a genus of 65 species that are broadly distributed throughout North America. Like other genera of Polemoniaceae, *Phlox* species have perfect flowers that are radially symmetrical, and each flower has a superior ovary with three chambers or locules. Placentation is axile – i.e. ovules initiate from placental tissue that lies along the central septum that subdivides the ovary into locules. In most species and in the outgroup *Microsteris gracilis*, exactly one ovule forms per locule (termed single-ovulate morphology) but six species have been described as producing more than one seed per locule in at least some fruits (termed multi-ovulate morphology) (Fig. 1; Wherry 1955). In species that produce the multi-ovulate phenotype, individuals display plastic variation for ovule number (and subsequent seed number) per chamber (Fig. S1). Thus, we define three broad classes of ovule packaging morphology: fixed single-ovulate (the ancestral condition), plastic single-ovulate (when a species that *can* produce more than one ovule per locule only produces one), and plastic multi-ovulate (two or more ovules per locule).

### Sample collection and tissue storage

We sampled from wild populations of multi-ovulate species *Phlox buckleyi*, *P. nana*, *P. nivalis*, *P. longifolia* and *P. roemeriana* and from wild populations of single-ovulate species *P. drummondii* and *M. gracilis* (Table S1). Additionally, we identified populations of *P. longifolia* that had a mixed phenotypic frequency of the multi-ovulate phenotype and others that were fixed for the single-ovulate phenotype. In our analysis, we treat *P. longifolia* individuals from populations that are fixed for the single-ovulate phenotype as “fixed single-ovulate”, categorically similar to *P. drummondii* and *M. gracilis*, and we treat individuals that originate from multi-ovulate populations of any phenotypic frequency as “plastic single-ovulate” (when they produce one ovule in at least one locule) or “plastic multi-ovulate” (when they produce two or more ovules in at least one locule).

Early bud tissue (Stage 1), ovaries from open flowers (Stage 2), and nearly mature fruits (Stage 3) were collected from multiple wild populations of each sampled species (Table S1; Fig. S1). Additionally, greenhouse-grown individuals of *P. drummondii* and *P. roemeriana* of single- or mixed-population ancestry were included. Early bud tissue and ovaries were immediately fixed in an FAA solution (50% ethanol, 4% formalin, and 5% glacial acetic acid) for at least 24 hours. Tissue was washed in 50% ethanol for at least 24 hours and then stored in 70% ethanol until histology or dissections were performed. Fruits were collected and stored in paper coin envelops to dry and dehisce. Voucher specimens of multiple representatives of each species were collected and deposited in the Harvard University Herbaria.

### Histological staining and measurements

To visualize ovary and ovule morphology early in bud development (Stage 1), we used standard histology and staining protocols (Hall et al. 2006). Bud tissue samples stored in 70% ethanol were further dehydrated to 100% ethanol and then embedded in Paraplast Plus (P3683, Sigma, Burlington, MA, USA). We sectioned samples at 8 µm on a Leica RM2235 microtome (Leica, Deer Park, IL, USA), and stained slides using a 0.1% Toluidine Blue O (T3260, Sigma, Burlington, MA, USA) solution for 3 minutes and mounted with a 1:1 ratio of Citrisolv (22-143-975, Fisher, Waltham, MA USA) to Permount (SP15-100, Fisher, Waltham, MA USA). Slides were imaged in the Harvard University Herbaria using a Zeiss Axioscope 7 (Zeiss, White Plains, NY, USA). To capture comparable developmental time points across samples, we defined the ovule initiation stage as the window of developmental time from the separation of the placental tissue from the ovary wall (i.e., the opening of the locule chamber) to the point at which initiated ovules begin to bend and become anatropous (Fig. 1, S1). This developmental window was found to be quite narrow across all species, although the exact bud width differed slightly: *M. gracilis*, 0.4-0.65 mm; *P. drummondii*, 0.6-1 mm; *P. roemeriana*, 0.7-1.1 mm; *P. longifolia*, 0.8-1.1 mm; *P. nana*, 1-1.5 mm; *P. buckleyi*, 0.8-1.1 mm; and *P. nivalis*, 0.6-0.95 mm.

### Ovary and fruit dissections

Fixed, unfertilized ovaries (Stage 2) were dissected and imaged at the Arnold Arboretum of Harvard University using a Zeiss Discovery version 12 dissecting microscope with an AxioCam 512 attachment (Zeiss, White Plains, NY, USA). A light micrograph was taken before dissection for ovary size quantification and after the removal of the ovary wall to quantify ovule size(s) (e.g., Fig. S1). If ovule number was asymmetric between locules of the same ovary, an image was taken to represent each unique ovule number per locule.

Dry fruits and seeds (Stage 3) were imaged in the Harvard University Herbaria using a Zeiss Discovery microscope (Zeiss, White Plains, NY, USA). For each individual specimen, we imaged three fruit pieces and three seeds per category (fixed single-ovulate morphology or plastic single-ovulate and plastic multi-ovulate).

### Measurements

We used ImageJ version 2.14.0 to measure morphological differences between ovule packaging categories at three stages: Stage 1, ovule initiation; Stage 2, open flowers; and Stage 3, mature fruits. For the Stage 1 histology sections, we measured ovary length, ovary width, ovule primordium diameter, surface length of the placental tissue, height of the placental tissue, petal length, and bud width (Figs. 2a-b, S1-2). In regard to the placental tissue, the extent of tissue proliferation at the base of the septum from which ovule primordia emerge, was remarkably variable within and between species. We define this tissue as the placenta, because the apical extent of the mass also corresponds to the uppermost position in which we observe ovule initiation along the central septum. To quantify the approximate size of the placental tissue and to capture the curvature obvious in some species, we use a ratio of the measured surface length of this proliferation (excluding ovule primordia) to the measured linear height of the proliferation from the base of the ovary to apex of the proliferation (Fig. 2a). For clarity, we use the term “placenta proliferation” to refer to this ratio. For each individual specimen, we measured up to two ovule primordia per locule and up to two locules per ovary. Each trait was generally measured on at least 3 serial sections to determine the maximum measurement and control for factors such as sample orientation. For each species, we examined at least five specimens.

**Fig. 2.**
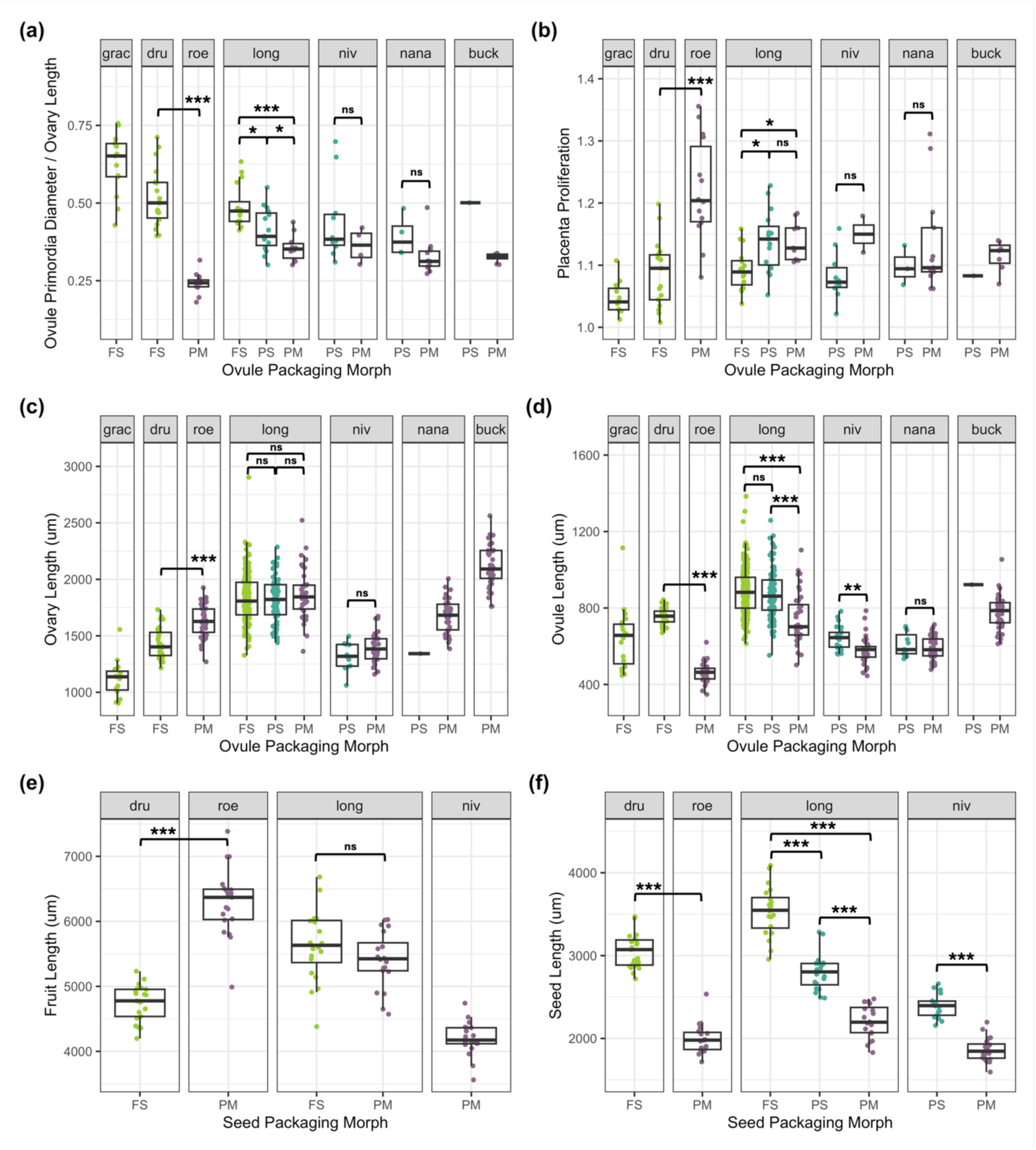
Morphological variation in gynoecium structures across three developmental stages. (a) Ratio of ovule primordia diameter to ovary length at the time of ovule initiation (Stage 1). (b) Ratio of placenta surface length to placenta height at the time of ovule initiation (Stage 1). (c) Ovary length and (d) ovule length sampled from open flowers (Stage 2). (e) Fruit length and (f) seed length sampled from collected mature fruits (Stage 3). T-tests were performed within species and between categories and between sister-species pair, *P. drummondii* and *P. roemeriana*. Significance is indicated by * : p < 0.05, **: p < 0.001, ***: p <0.0001, ns: p > 0.05. Species names are abbreviated as: *M. gracilis* = grac, *P. drummondii* = dru, *P. roemeriana* = roe, *P. longifolia* = long, *P. nivalis* = niv, *P. nana* = nana, *P. buckleyi* = buck. Morph categories are abbreviated as: fixed single = FS, plastic single = PS, plastic multi = PM.

For the Stage 2 dissected ovaries, we measured ovary length, ovary width, ovule length, and ovule width (Figs. 2c-d, S1f-g, S3). For the Stage 3 fruits, we measured fruit length, fruit width, seed length, and seed width (Figs. 2e-f, S4, S1h-j).

### Statistical Analyses

To account for variation in measurements across Stage 1 histological sections of the ovaries, we used the maximum measurements derived from three successive sections for ovary length, ovary width, each ovule primordium diameter, and petal length. Similarly, we used the maximum ratio between placenta surface length and placenta height for subsequent analyses. If multiple ovules were measured in a given locule, we averaged the maximum measurements for each ovule primordium. Measurements taken from each locule were treated as separate observations for the mixed effects models described below, controlling for ovary and individual. We performed pairwise correlations with Bonferroni correction for multiple comparisons between ovary size metrics and petal length, including all Stage 1 observations (by ovary, not locule) irrespective of species or ovule packaging morphology (Table S2). Because petal length is highly correlated with ovary length (r = 0.8410; Table S2, Fig. S2f) and width (r = 0.8436), we used petal length to control for trait variance that is due to continuous developmental time across the defined discrete ovule initiation stage

For Stage 2, we treated ovule measurements from different locules as separate observations only if the number of ovules per locule was asymmetric (i.e. different numbers of ovules in different locules). In Stage 3, we separated measurements of single- and multi-ovulate morphology seeds, if present in a single individual. In both cases, if measurements were separated, we controlled for the ovary or individual, respectively.

All analyses were performed in R version 4.5.1 (R Core Team 2025). We performed linear mixed effects (LME) models using the lme4 package (Bates et al. 2015) to test our hypotheses. First, we fit an LME model with ovule primordia diameter as the dependent variable; an interaction between ovary length and ovule packaging morphology; species and petal length as fixed factors; and individual and bud tissue sample as a nested random factor (Table 1). After failing to detect a significant effect of the interaction term on the dependent variable, we conducted two simplified follow-up analyses. First, we collapsed the “plastic single-ovulate” and “plastic multi-ovulate” categories into a single group and reclassified samples according to genotype (i.e. fixed single-ovulate genotype or multi-ovulate genotype) (Table 1). We used the interactions package and an emtrends post hoc analysis to test whether the estimated slopes of the linear models were significantly different from zero and from one another (Tables S3, S4; Lenth 2017; Long 2019). Second, we ran a simplified version of the original model that retained the original ovule packaging categories but excluded the interaction term (Table 1). We then used an emmeans posthoc test with a Tukey method correction for multiple comparisons to test for pairwise significance between fixed single-, plastic single-, and plastic multi-ovulate categories in our model (Table S5; Lenth 2017). Next, we performed unpaired t-tests between morph categories of individuals of the same species and between the sister species pair *P. drummondii* and *P. roemeriana* to test for significant differences in ovule primordium diameter relative to ovary length (Fig. 2a).

**Table 1.**
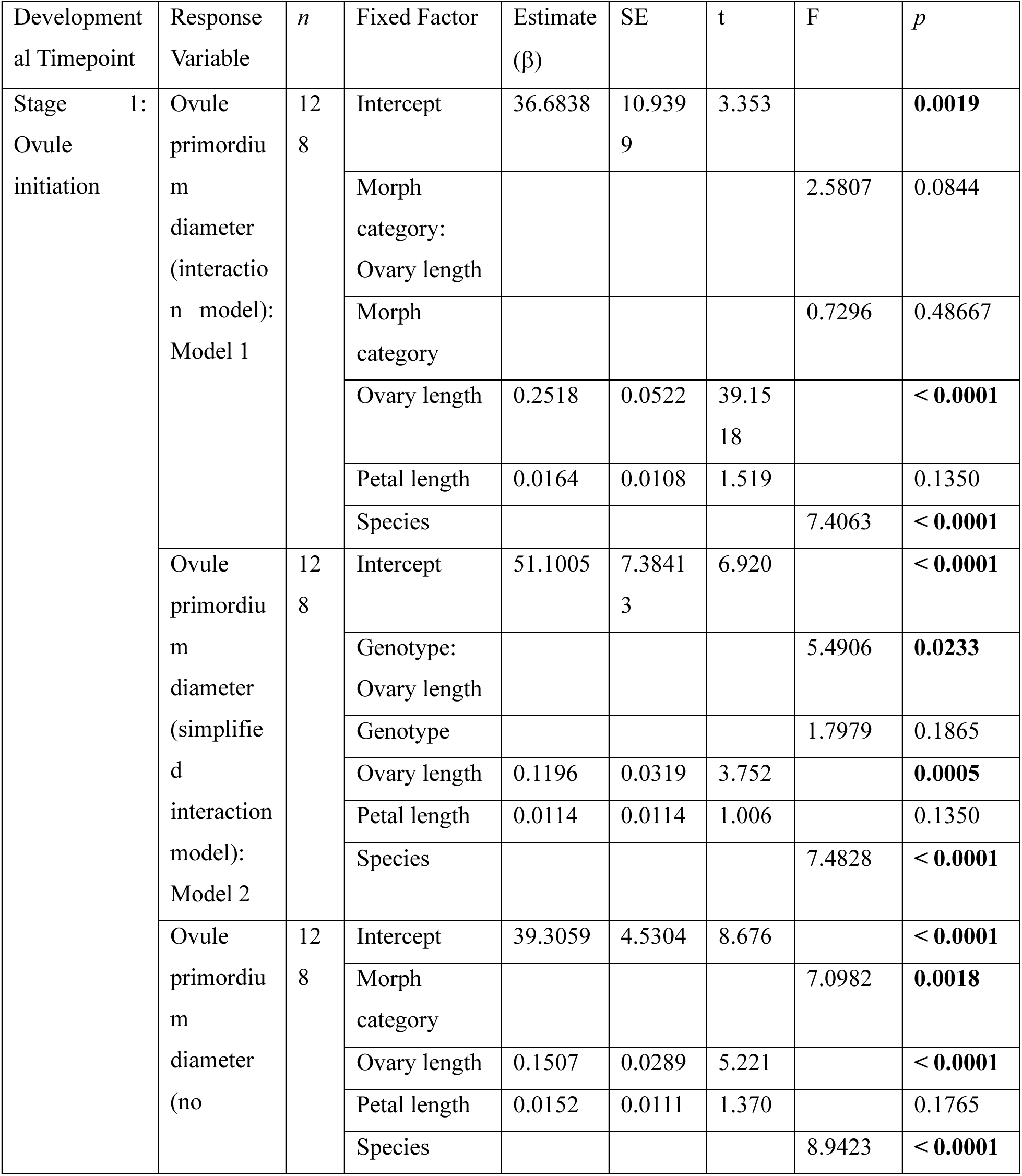
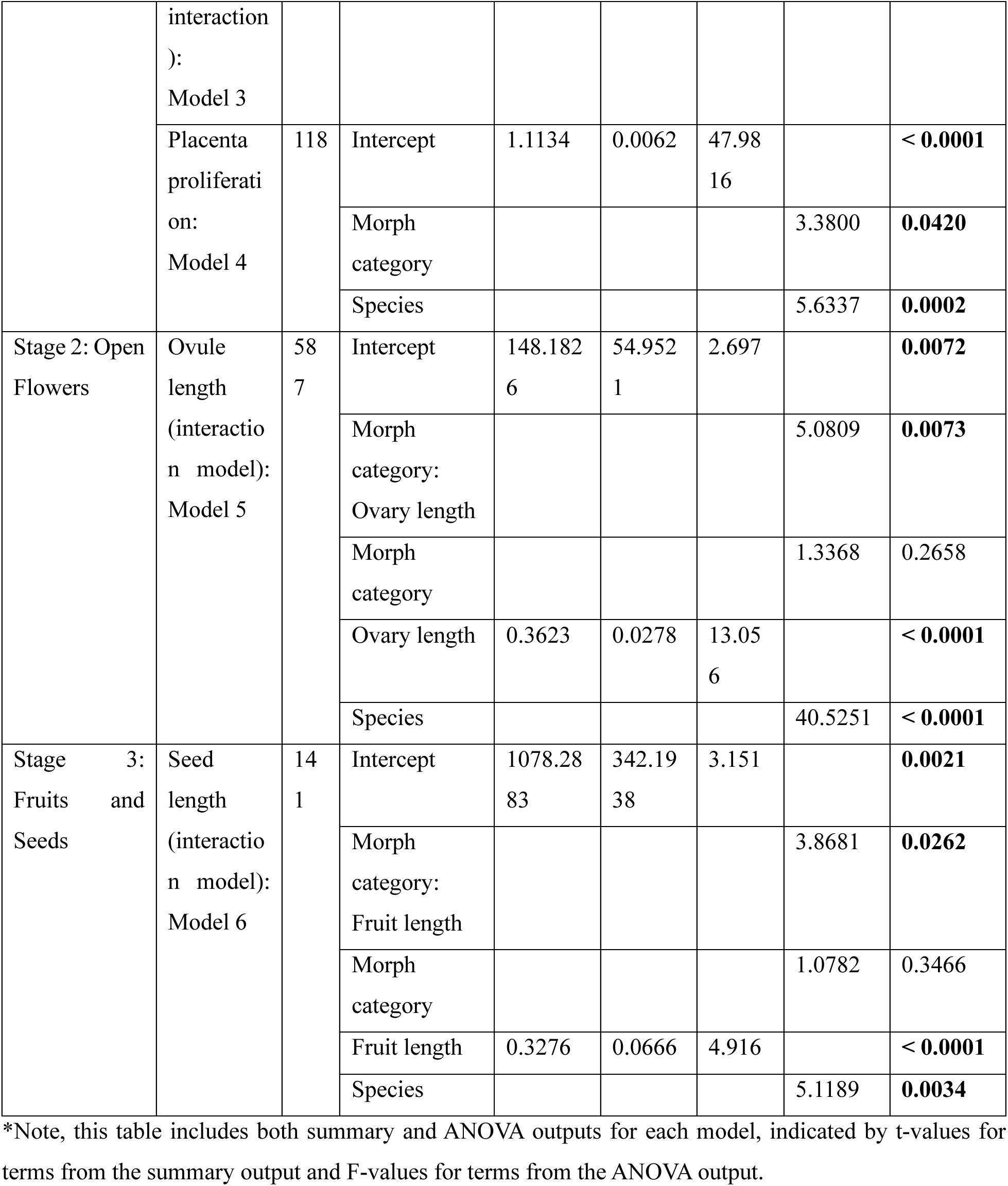
Mixed-effects models to test for changes in allometric relationships across developmental time.

To test for differences in placenta proliferation between ovule packaging morphs, we use an LME model in which placenta proliferation is the dependent variable; ovule packaging morph and species are treated as fixed factors; and individual and bud tissue sample are a nested random factor. We also used emmeans with a Tukey correction for multiple comparisons as a posthoc test of significance within the model and pairwise t-tests within and between species as before (Fig. 2b, Table S5; (Lenth 2017).

We took a similar approach to understanding the role of ovule packaging morphology on ovule size and the relationship between ovule size and ovary size in open flowers (Stage 2). To test for a change in allometry between ovules and ovaries from open flowers, we fit a model with ovule length as the dependent variable; an interaction between ovary length and ovule packaging morph fixed factor; species as an independent fixed factor; and population and individual as a nested random factor. We evaluated the slope of ovule length by ovary length across ovule morph categories using pairwise β-coefficients and the sim_slopes function (Table S2; (Long 2019). We also used the emtrends posthoc analysis to test whether the slope of the relationship between ovary length and ovule length differed between treatments (Table S3; (Lenth 2017). We used unpaired t-tests for comparisons of ovule length and width and ovary length and width within species and between sister species *P. drummondii* and *P. roemeriana* (Fig. 2c,d).

Finally, in Stage 3, to test for a difference in slope between seed length and fruit length across ovule packaging morphs, we used an LME model with seed length as the dependent variable; an interaction between fruit length and ovule packaging morph; species as an independent fixed factor; and population and individual as a nested random factor (Table 1). We compared pairwise β-coefficients between interaction terms. We followed with sim_slopes and emtrends posthoc tests as before (Tables S3-S4). We used unpaired t-tests for all pairwise comparisons within species or between species pair *P. drummondii* and *P. roemeriana* (Fig. 2e,f).

## RESULTS

### Stage 1: Ovule initiation & ovary development

Surprisingly, previous studies of variation in seed number per fruit have generally not focused on the critical first step in seed development, the initiation of the ovules themselves. This means that we lack important information on the relationship between ovary and ovule primordium size, and how these relate to ovule number. Obviously, ovary length and width and ovule primordium diameter vary within and between species (Fig. S2a-c). However, this variation may be confounded by several factors, including continuous developmental time across ovule initiation and floral size variation within the inflorescence. To control for variation due to developmental stage and inflorescence position, we examined other landmarks of floral morphology. Petal length is highly correlated with ovary length and width when considering all bud samples included in Stage 1 (Fig. S2f; Table S2; r = 0.8410 and r = 0.8436, respectively), so we included petal length as a fixed effect in the LME model to account for this variation. Further, we present relative ovule primordium size as a ratio of ovule primordium diameter to ovary length, which corrects for size variation in both structures and focuses our analysis on the allocation to ovule size relative to ovary size (Fig. 2a). While we find the same qualitative pattern of allometric relationships between ovary size and ovule size for all three ovule packaging categories at Stage 1 compared to Stages 2 and 3 (Fig. S2d), we failed to find a significant interaction term between ovary length and morph category in Model 1 (Table 1). Thus, we used two simplified models instead. With Model 2, we find that the slope of the relationship between ovary length and ovule primordium diameter is significantly reduced in multi-ovulate genotypes compared to single-ovulate genotypes (Table 1; F = 5.4906, p = 0.0233). We find that the slope of the relationship between ovary length and ovule primordium diameter in single-ovulate and multi-ovulate genotypes are both significantly different from zero (Table S3; single-ovulate genotype: β = 0.2588, p < 0.0001, multi-ovulate genotype: β = 0.0319, p = 0.0005) and from each other (Table S4; Fig. 1.4e; Δβ = - 0.1390, p = 0.0233). Second, we ran an LME model that controlled for variation explained by ovary length, petal length, and species without an interaction between ovary length and morph category (Table 1). In this model, we found that ovule primordium diameter significantly differed across morph categories and by species and that ovary length was a significant predictor of ovule primordium diameter (Table 1). A posthoc test revealed that fixed single-and plastic single-ovulate ovaries had significantly larger ovule primordia than plastic multi-ovulate ovaries (Table S5; ΔEMM = 15.36 um, p = 0.0053 and ΔEMM = 7.16 um, p =0.0111, respectively), but there was no difference in ovule primordium diameter between fixed single- and plastic single-ovulate ovaries (ΔEMM = 8.20 um, p = 0.1565). Most paired comparisons within species and between the sister species *P. drummondii* and *P. roemeriana* show that relative ovule primordia size is smaller in multi-ovulate locules compared to single-ovulate locules (Fig. 2a).

Using an LME model, we found that placental proliferation varies by morph category and by species (Table 1, Fig. 2b). Using a posthoc test, we find that plastic multi-ovulate ovaries have greater placental proliferation compared to fixed single-ovulate ovaries (Table S5; ΔEMM = 0.0563, p = 0.0334). Similar to our findings on relative ovule primordia size, we observe an intermediate morphology for plastic single-ovulate ovaries in comparison to fixed single- and plastic multi-ovulate ovaries, but neither comparison is significant (Table S5; ΔEMM = 0.0358, p = 0.1839 and ΔEMM = -0.0205, p = 0.4358, respectfully). Notably, placental proliferation is greatest in the most extreme multi-ovulate phenotype found in *P. roemeriana* (Fig. 2b). Placental proliferation in *P. roemeriana* is 12% greater than its single-ovulate sister species *P. drummondii* on average.

### Stage 2: Pre-fertilization ovules & ovaries

We ran an LME model with an interaction term between morph category and ovary length to evaluate the slope of the relationship between ovule length and ovary length across categories (Table 1). We find a significant interaction between ovary length and morph category (Table 1; F = 5.0809, p = 0.0073) All three morph categories have positive slopes that are significantly different from zero (Table S3; fixed single: β = 0.3623, p < 0.0001, plastic single: β = 0.2774, p < 0.0001, plastic multi: β = 0.2386, p < 0.0001). Using an emtrends posthoc test with a Tukey method multiple comparisons correction, we find that the slope of the relationship between ovary size and ovule size is significantly reduced in plastic multi-ovulate individuals relative to fixed single-ovulate (Table S4; Fig. S3c; Δβ = -0.1237, p = 0.0096). Similar to Stage 1, the plastic single-ovulate group has an intermediate slope between the fixed single- and plastic multi-ovulate groups; however, we do not find a significant difference in slope between the plastic single-ovulate morph category and either the fixed single- or plastic multi-ovulate groups (Table S4; Δβ = 0.0849, p = 0.1070 and Δβ = -0.0388, p = 0.1359, respectfully).

Ovary size in pre-fertilization flowers is remarkably variable across species but shows no clear relationship with ovule packaging strategy (Figs. 2c). Multi-ovulate species *P. roemeriana* has significantly larger ovaries than single-ovulate species *P. drummondii,* but we find no difference in ovary size between morph categories within any species (Fig. 2c). Conversely, ovule size is significantly smaller in multi-than single-ovulate ovaries in all paired comparisons except *P. nana* (Figs. 2d). Interestingly, in *P. longifolia* we find no difference in ovule size between fixed single- and plastic single-ovulate ovaries, indicating that the differences in relative ovule primordia size described in Stage 1 do not result in ovule size differences at the time of floral opening.

### Stage 3: Mature fruits & seeds

Using an LME model with an interaction between morph category and fruit length, we again find a significant interaction term (Table 1; F = 3.8681, p = 0.0262), and a posthoc test with a Tukey method correction for multiple comparisons shows a significant reduction in slope from the fixed single-ovulate group to the plastic multi-ovulate, with an intermediate allometry for the plastic single-ovulate group that is not significantly different from either the fixed single-ovulate group or the plastic multi-ovulate group (Fig. S4c, Table S4; fixed single – plastic multi: Δβ = 0.2210, p = 0.0272; fixed single – plastic single: Δβ = 0.1663, p = 0.501; plastic single-plastic multi: Δβ = 0.0547, p = 0.3957). The positive slopes of all three categories are significantly different from zero (Fig. S4c, Table S3; fixed single: β = 0.3276, p < 0.0001, plastic single: β = 0.1613, p = 0.0067, plastic multi: β = 0.1066, p = 0.0429).

Similar to ovary size, we see notable variation in fruit size that is not readily attributed to ovule packaging strategy (Fig. 2e). However, *P. roemeriana* does have the largest fruits (Fig. 2e), which likely facilitates the production of 30 seeds per fruit, or (rarely) more. We also find that across all pairwise comparisons, multi-ovulate seeds are significantly smaller than single-ovulate seeds (Fig. 2f), consistent with the hypothesis of a fundamental tradeoff between seed size and number. Further, the significant difference in seed size between fixed single- and plastic single-ovulate seeds of *P. longifolia* suggests a tradeoff between the fixed single-ovulate strategy and the plastically variable multi-ovulate strategy (Fig. 2f).

## DISCUSSION

Here we demonstrate strong empirical evidence that the transition from a single-ovulate to a multi-ovulate strategy is accompanied by a shift in the allometric relationship between ovule size and ovary size that begins at ovule initiation and continues through seed development, resulting in a tradeoff between maternal reproductive fitness components: seed size and number. Specifically, we observe two distinct forms of developmental plasticity. In the ancestral, single-ovulate condition, larger ovaries result in larger ovule primordia but never more than one ovule; while in the derived multi-ovulate condition, the ovules tend to be smaller relative to the ovary, allowing at least two ovules per locule, and increased ovary size can result in the production of additional ovules (Fig. 2, S2d, S3c, S4c). Further, we find that the plastic single-ovulate phenotype produced by multi-ovulate individuals results in an intermediate allometry and intermediate allocation pattern across all developmental stages when compared to fixed single-ovulate and plastic multi-ovulate ovaries, indicating that even when multi-ovulate ovaries initiate a single ovule, they are not equivalent to the ancestral condition (Fig. 2a and f, S2d, S3c, S4c). Finally, we report increased placental proliferation in multi-ovulate ovaries compared to fixed single-ovulate ovaries. We observe this placental proliferation to be most pronounced in *P. roemeriana,* likely contributing to the extreme multi-ovulate phenotype present in this species (Fig. 2b).

Studies of the genetic basis of seed number per fruit have focused almost exclusively on *Arabidopsis thaliana* (reviewed in Barro-Trastoy et al. 2020; Cucinotta et al. 2020). From a developmental genetic perspective, we know that ovule initiation is controlled by a combination of hormone pathways, particularly auxin, cytokinin, gibberellins and brassinosteroids. Auxin is perhaps the most important of these as its transport and response is essential to the specification of ovule primordia (reviewed Qadir et al., 2021). The loci underlying all of these pathways tend to be highly pleiotropic and reveal a strong correlation between ovary size, placental size and ovule number in this system (Cucinotta et al. 2020), which is also observed in genome-wide association studies of variation in seed number per silique in both *Arabidopsis thaliana* and *Brassica napus* (Yuan & Kessler 2019; Jiang et al., 2020; Qadir et al., 2022). In mutant backgrounds, disrupting relevant hormonal regulation can increase or decrease ovary size (and placenta surface area), resulting in a change in ovule primordia number, while the primordium size appears to remain the same. Even mutants that dramatically increase spacing between ovules do not alter ovule size (Galbiati et al. 2013). Based on this work, one model for the transition from single-ovulate to multi-ovulate ovaries would be a simple increase in ovary size without modification of ovule size. However, that does not appear to be the case in *Phlox*. Thus, our current models for understanding ovule number variation may be missing an important source of variation underlying morphological complexity, and discrete shifts in ovule packaging strategy (as opposed to quantitative differences in ovule number in multi-ovulate lineages) may require a different set of genetic and developmental mechanisms.

We also demonstrate that ovule primordia size varies plastically, not only with ovary size, but also depending on the presence or absence of other ovule primordia in the locule. However, the differences in relative ovule primordia size in plastic single-ovulate compared to fixed single-ovulate ovaries provides evidence that the regulation of ovule primordia size is at least partially decoupled from ovary size in the derived multi-ovulate state. This is particularly evident in *P. longifolia*. In this species, we observed that some populations across the species range are fixed for the single-ovulate phenotype, while other populations have variable frequencies of both morphologies (Bickner, unpublished data). Notably, the striking difference in seed size between fixed single-ovulate and plastic single-ovulate *P. longifolia* individuals (Fig. 2f) suggests that there is a potential cost to the multi-ovulate strategy: single-ovulate seeds from multi-ovulate plants are smaller. However, further studies are necessary to disentangle the relative roles of genetic variation and plasticity in producing the observed phenotypic distributions of this species.

Finally, the transition to a multi-ovulate strategy in *P. roemeriana* is particularly extreme. Consistent with our findings in other multi-ovulate species, we observe that reduced ovule primordia size is an important contributor to ovule packaging strategy divergence (Figs. 2a, S2b) as well as ovary size (Fig. S2a). However, we also observed a unique proliferation of the placental tissue that increases the surface area for ovule primordia formation independent of ovary size (Figs. S1, 2b). Modification of placental morphology has been shown to underlie ovule number variation in many lineages (Ronse De Craene 2021; Shivaprakash and Bawa 2022) but Phlox is particularly useful in exhibiting shifts in placenta proliferation across a microevolutionary timescale, differentiating closely-related sister species. Thus, future comparative work describing the genetic regulation of placental proliferation in this species pair could provide novel insight into mechanisms of reproductive strategy divergence that are more often obscured by deep evolutionary time.

Collectively, our findings suggest a repeated pattern of developmental evolution in independently derived multi-ovulate lineages, including how plasticity produces ovule number variation. This study highlights the potential of the *Phlox* genus as a model to understand the evolution of ovule packaging strategies, the restructuring of fundamental allometric relationships, and, ultimately, the genetic basis of tradeoffs in maternal fitness.

## ACKNOWLEDGEMENTS

We are grateful to Y. Yildirim for assistance with histological sectioning; S. Pesce, J. Nelson, L. Lee, M.J. Epps, and E. Roalson for assistance in the field; and A. Serrato-Capuchina and S. Chaturvedi for supplying some fruit and seed samples. We thank the Bureau of Land Management; the Washington Department of Natural Resources; the South Carolina Department of Natural Resources; the Uinta, Wasatch, Cache, Cibola, George Washington and Jefferson National Forests; Pagosa Springs Veterans Park (CO); Tonasket School District (WA); Forty Acre Rock Heritage Preserve/Wildlife Management Area; the Carolina Sandhills National Wildlife Refuge; and the Department of Energy at Savannah River Site for permitting and land use permission. We also thank the Commonwealth of Virginia Department of Conservation and Recreation for occurrence data of *P. buckleyi*. We acknowledge the Arnold Arboretum of Harvard University and the Harvard University Herbaria for use of research facilities for imaging. T. Kellogg, P. Stevens, D. Faccini, R. Hopkins, D. Haig, and the Kramer Lab provided helpful discussions and comments on previous versions of this manuscript. This work was funded in part by awards through the Harvard University Herbaria and the Harvard Graduate Student Council to B.M.B. and from internal funding through the Harvard University Department of Organismic and Evolutionary Biology to B.M.B.

## Notes

https://doi.org/10.5281/zenodo.18165895

### Competing Interest Statement

The authors have declared no competing interest.

