## Supplementary Figures and Tables for "A shift in developmental allometry underlies the transition to a multi-ovulate strategy from a single-ovulate ancestral state in *Phlox* (Polemoniaceae)"

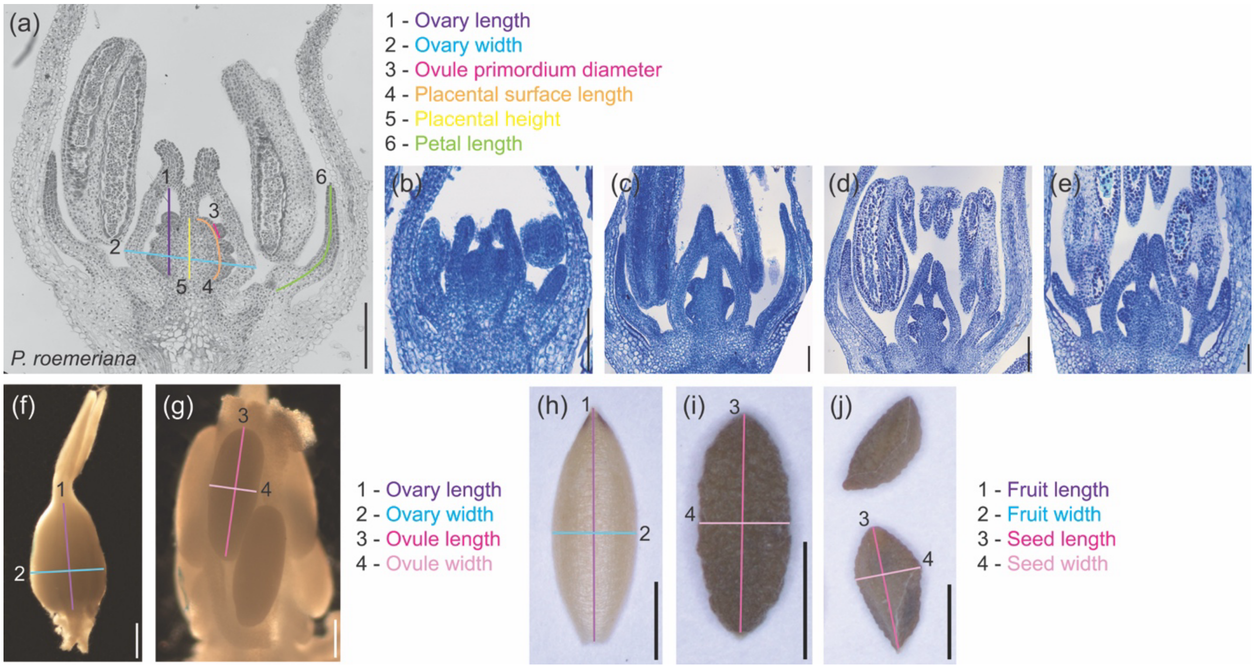


**Fig. S1**. All measurements taken with example histological sections of remaining selected taxa. (a) Grayscale *P. roemeriana* section with all measurements indicated: 1) Ovary length (purple), 2) Ovary width (blue), 3) Ovule primordium diameter (pink), 4) Placental surface length (orange), 5) Placental height (yellow), 6) Petal length (green). (b) *M. gracilis* section (single-ovulate). (c) *P. nana* section (asymmetric: left locule is bi-ovulate, right locule is plastic single-ovulate). (d) *P. longifolia* section (bi-ovulate). (e) *P. nivalis* section (bi-ovulate). (f) *P. roemeriana* fixed carpel with ovary length (1/purple) and ovary width (2/blue). (g) *P. longifolia* fixed, dissected ovary with ovule length (3/dark pink) and ovule width (4/light pink). (h) *P. longifolia* fruit valve with fruit length (1/purple) and fruit width (2/blue). (i) fixed single-ovulate *P. longifolia* seed with seed length (3/dark pink) and seed width (4/light pink). (j) multi-ovulate *P. longifolia* with seed length (3/dark pink) and seed width (4/light pink). Size bars in (a, d-e, g) = 200 µm; (b-c) = 100 µm; (f) = 500 µm; (h-j) = 2000 µm.

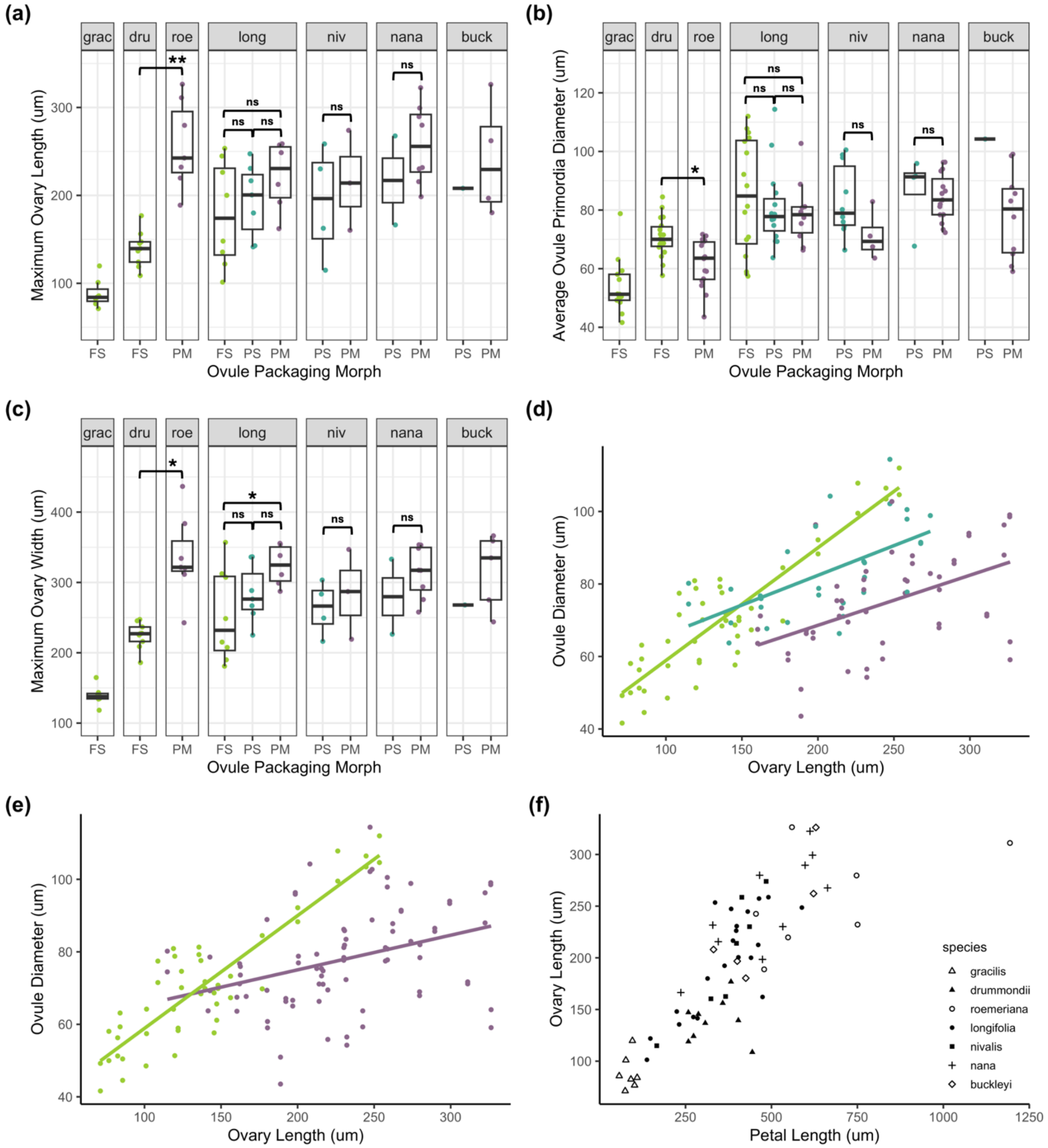


**Fig. S2**. Stage 1 raw data (a-c) and relationships between ovary length vs. petal length (d) and ovary length vs. ovule diameter (e). (a) Ovary length by ovule packaging morphology and species. (b) Ovule primordia diameter by ovule packaging morphology and species. (c) Ovary width by ovule packaging morphology and species. (d) Linear models of the relationship between ovary length and ovule primordia diameter for fixed single-ovulate (green), plastic single-ovulate (blue), and multi-ovulate (purple) morph categories. (e) Simplified linear models of the relationship between ovary length and ovule primordium diameter for fixed single-ovulate (green) and all samples from multi-ovulate genotype individuals (purple). (f) Positive correlation between petal length and ovary length at the time of ovule initiation. Significance is indicated by * :  $p < 0.05$, **: $p < 0.001$, ***: $p < 0.0001$. Species names are abbreviated as: *M. gracilis* = grac, *P. drummondii* = dru, *P. roemeriana* = roe, *P. longifolia* = long, *P. nivalis* = niv, *P. nana* = nana, *P. buckleyi* = buck. Morph categories are abbreviated as: fixed single = FS, plastic single = PS, plastic multi = PM.

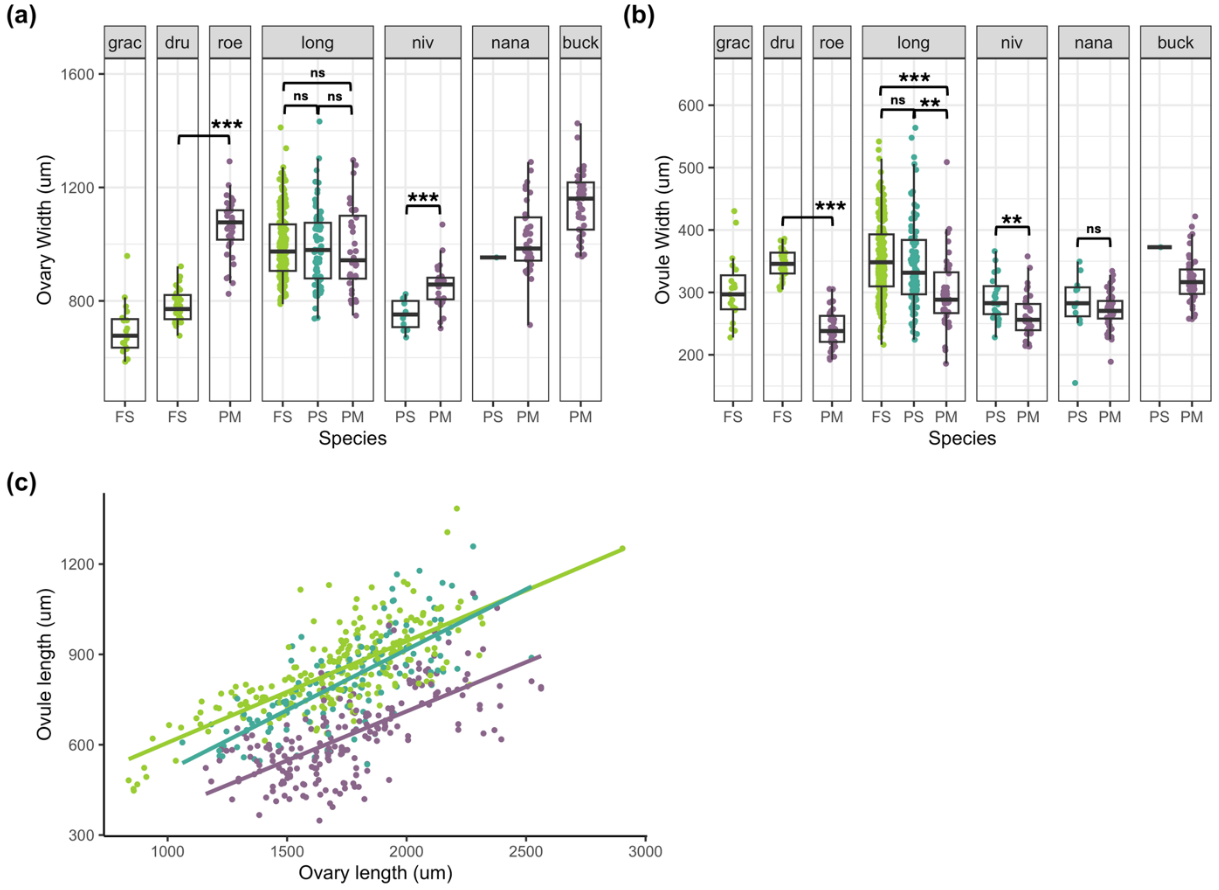


**Fig. S3**. Stage 2 data (a-b) and relationships between ovary length, petal length and ovule diameter (c). (a) Ovary width by ovule packaging morphology and species. (b) Ovule width by ovule packaging morphology and species. (c) Linear models of the relationship between ovary length and ovule length for fixed single-ovulate (green), plastic single-ovulate (blue), and multi-ovulate (purple) morph categories. Significance is indicated by * : p < 0.05, **: p < 0.001, ***: p <0.0001. Species names are abbreviated as: *M. gracilis* = grac, *P. drummondii* = dru, *P. roemeriana* = roe, *P. longifolia* = long, *P. nivalis* = niv, *P. nana* = nana, *P. buckleyi* = buck. Morph categories are abbreviated as: fixed single = FS, plastic single = PS, plastic multi = PM.

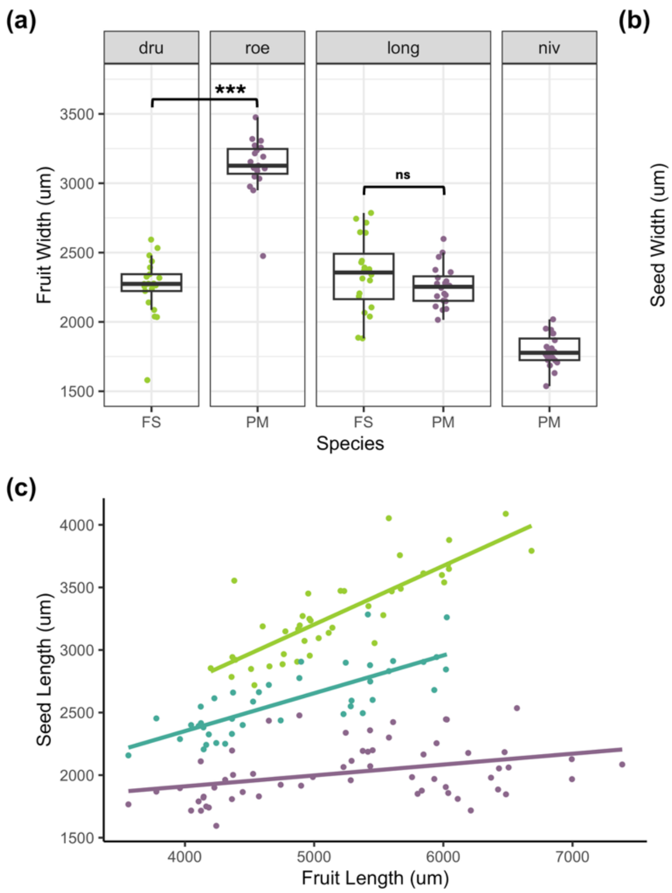


**Fig. S4**. Stage 3 data (a-c) and relationships between ovary length, petal length and ovule diameter (d-e). (a) Fruit width by ovule packaging morphology and species. (b) Seed width by ovule packaging morphology and species. (c) Linear models of the relationship between fruit length and seed length for fixed single-ovulate (green), plastic single-ovulate (blue), and multi-ovulate (purple) morph categories. Significance is indicated by * : $p < 0.05$, **: $p < 0.001$, ***: $p < 0.0001$. Species names are abbreviated as: *M. gracilis* = grac, *P. drummondii* = dru, *P. roemeriana* = roe, *P. longifolia* = long, *P. nivalis* = niv, *P. nana* = nana, *P. buckleyi* = buck. Morph categories are abbreviated as: fixed single = FS, plastic single = PS, plastic multi = PM.

**Table S1**. Source of tissue sample collection

| Population | Species | Latitude | Longitude |
|---|---|---|---|
| L006G | *Microsteris gracilis* | 46.7862 | -117.4117 |
| L023G | | 48.22937 | -119.76416 |
| B002 | *Phlox buckleyi* | | |
| B003 | | | |
| B004 | | | |
| B011 | | | |
| 680A | *P. drummondii* | 29.746617 | -97.4097 |
| 656A | | 30.0852333 | -96.9161667 |
| 660 | | 30.1803833 | -97.4207167 |
| 662 | | 30.0621514 | -97.3502951 |
| 667 | | 29.528817 | -96.441233 |
| 669 | | 29.965443 | -96.885429 |
| 676 | | 29.4689024 | -97.8737457 |
| 677 | | 28.842167 | -97.845583 |
| 679 | | 29.367483 | -97.566617 |
| 690 | | 30.880867 | -98.653667 |
| 702 | | 30.901115 | -99.288657 |
| 746 | | 29.218909 | -98.972419 |
| Family 8 | | 29.52881 | -96.44123 |
| | | 29.71848 | -96.8887 |
| Family 10 | | 32.12872 | -98.31181 |
| L015/L184 | *P. longifolia* | 47.52316 | -120.0464 |
| L023 | | 48.22937 | -119.76416 |
| L031/L178 | | 48.70271 | -119.43067 |
| L039 | | 47.66071 | -117.68799 |
| L102 | | 37.76884 | -103.62091 |
| L103 | | 39.64058 | -106.57223 |
| L104 | | 39.46949 | -107.26382 |
| L105 | | 39.11943 | -108.31852 |
| L106 | | 39.25352 | -108.9304 |
| L107 | | 39.45837 | -110.48414 |
| L108 | | 40.38014 | -111.98868 |
| L109 | | 40.38281 | -112.02176 |
| L110 | | 40.23655 | -112.19878 |
| L111 | | 40.35146 | -112.30748 |
| L112 | | 40.51157 | -112.31239 |
| L113 | | 41.84682 | -112.06033 |
| L114 | | 41.8249 | -112.07831 |
| L115 | | 41.21578 | -112.47325 |
| L116 | | 40.22476 | -111.62283 |
| L117 | | 40.04611 | -111.47633 |
| L118 | | 39.9751 | -111.69233 |
| L119 | | 40.21253 | -110.85873 |
| L120 | | 40.20648 | -110.38916 |
| L121 | | 40.05521 | -111.80652 |

| | | | |
|---|---|---|---|
| L122 | | 39.43106 | -110.86911 |
| L123 | | 38.42497 | -109.43301 |
| L124 | | 38.27979 | -109.2967 |
| L125 | | 37.83307 | -109.10084 |
| L126 | | 37.99107 | -108.93881 |
| L127 | | 38.09541 | -108.53105 |
| L128 | | 38.0021 | -108.03698 |
| L129 | | 36.87266 | -107.89116 |
| L130 | | 36.53642 | -107.87769 |
| L131 | | 37.22554 | -107.70178 |
| L132 | | 37.24746 | -107.09354 |
| L136 | | 37.652 | -109.51762 |
| L138 | | 38.48328 | -111.54905 |
| L145 | | 38.63569 | -111.84346 |
| L148 | | 39.02515 | -114.6715 |
| L153 | | 39.44288 | -115.93066 |
| L154 | | 39.45681 | -116.72645 |
| L155 | | 39.49255 | -117.05932 |
| L161 | | 40.93508 | -115.87991 |
| L183 | | 47.84864 | -119.96907 |
| L196 | | 43.49528 | -113.55219 |
| L198 | *P. nana* | 35.03739 | -106.3455 |
| L199 | | 35.16258 | -106.36997 |
| L200 | | 35.61999 | -105.92082 |
| L201 | | 35.57036 | -105.75152 |
| N003 | *P. nivalis* | 34.6599 | -80.51678 |
| N015 | | | |
| N016 | | 34.14511 | -80.8896 |
| N017 | | 34.56253801 | -80.19320397 |
| 681 | *P. roemeriana* | 29.9346 | -98.2605167 |
| 682 | | 29.7731 | -98.2814833 |
| 687 | | 31.1708000 | -100.5119000 |
| 689/720 | | 30.9591720 | -100.4570040 |
| 693 | | 30.480633 | -98.198200 |
| 700 | | 29.582519 | -98.693396 |
| 717 | | 30.800277 | -99.834818 |
| 728 | | 29.961564 | -99.580216 |
| Family 8 | | 29.961564 | -99.580216 |
| | | 30.493663 | -98.18217 |

Populations are identified as either fixed for the single-ovulate morphology (green) or with the multi-ovulate phenotype present at any frequency (light purple). GPS data for *P. buckleyi* populations and N015 population of *P. nivalis* are not available for publication due to permit and/or land use restrictions with the Commonwealth of Virginia Department of Conservation and Recreation and the United States Department of Energy at Savannah River Site. Contact Bickner for inquiries on population data for these collection sites.

**Table S2**. Pairwise trait correlations (Spearman) with Bonferroni multiple comparison correction. Correlation coefficients are above the diagonal and associated *p*-values are given below the diagonal.

| | | |
|---|---|---|
| Maximum ovary length (um) | r = 0.9494 | r = 0.8410 |
| p < 0.0001 | Maximum ovary width (um) | r = 0.8436 |
| p < 0.0001 | p < 0.0001 | Maximum petal length (um) |

**Table S3**. Sim-slopes posthoc analysis to test whether slopes are significantly different from zero

| Stage | Model | Ovule packaging category | Estimate (β) | SE | t-ratio | *p* |
|---|---|---|---|---|---|---|
| Stage 1 | Model 2 | Single-ovulate | 0.2588 | 0.0566 | 4.5762 | **<0.0001** |
| | | Multi-ovulate | 0.1196 | 0.0319 | 3.7524 | **0.0005** |
| Stage 2 | Model 5 | FS | 0.3623 | 0.0277 | 13.0559 | **<0.0001** |
| | | PS | 0.2774 | 0.0314 | 8.8370 | **<0.0001** |
| | | PM | 0.2386 | 0.0317 | 7.5326 | **<0.0001** |
| Stage 3 | Model 6 | FS | 0.3276 | 0.0666 | 4.9163 | **<0.0001** |
| | | PS | 0.1613 | 0.0585 | 2.7599 | **0.0067** |
| | | PM | 0.1066 | 0.0519 | 2.0533 | **0.0429** |

*Note: FS = Fixed single-ovulate, PS = Plastic single-ovulate, PM = Plastic multi-ovulate

**Table S4**. Emtrends posthoc tests for pairwise comparisons between ovule packaging morphs of the slope of the relationship between ovule length and ovary length

| Stage | Model (see Fig. 1.3) | Paired comparison | Estimated difference in slope (β) | SE | df | t-ratio | *p* |
|---|---|---|---|---|---|---|---|
| Stage 1 | Model 2 | Single-ovulate – Multi-ovulate | 0.139 | 0.0594 | 47.6 | 2.343 | **0.0233** |
| Stage 2 | Model 5 | FS - PM | 0.1237 | 0.0421 | 544.1 | 2.937 | **0.0096** |
| | | FS – PS | 0.0849 | 0.0419 | 539.2 | 2.026 | 0.1070 |
| | | PM – PS | -0.0388 | 0.0201 | 88.7 | -1.931 | 0.1359 |
| Stage 3 | Model 6 | FS - PM | 0.2210 | 0.0845 | 109.7 | 2.615 | **0.0272** |
| | | FS – PS | 0.1663 | 0.0886 | 118.5 | 1.876 | 0.1501 |
| | | PM – PS | -0.0547 | 0.0416 | 42.5 | -1.313 | 0.3957 |

*Note: FS = Fixed single-ovulate, PS = Plastic single-ovulate, PM = Plastic multi-ovulate

**Table S5**. Emmeans posthoc tests for pairwise comparisons between ovule packaging morphs

| Stage | Model (see Fig. 1.3) | Paired comparison | Estimated difference in marginal means | SE | df | t-ratio | *p* |
|---|---|---|---|---|---|---|---|
| Stage 1 | Model 3 | FS - PM | 15.36 um | 4.65 | 44.3 | 3.301 | **0.0053** |
| | | FS – PS | 8.20 um | 4.35 | 39.5 | 1.885 | 0.1565 |
| | | PM – PS | -7.16 um | 2.44 | 110.0 | -2.940 | **0.0111** |
| Stage 1 | Model 4 | FS - PM | -0.0563 | 0.0217 | 42.8 | -2.600 | **0.0334** |
| | | FS – PS | -0.0358 | 0.0199 | 38.0 | -1.798 | 0.1839 |
| | | PM – PS | 0.0205 | 0.0166 | 101.9 | 1.234 | 0.4358 |

*Note: FS = Fixed single-ovulate, PS = Plastic single-ovulate, PM = Plastic multi-ovulate
